# Design and development of a SARS and MERS Combination Vaccine

**DOI:** 10.1101/2025.10.27.683653

**Authors:** Antonio Casal, Mohamed D. Yousif, Samuel Ellis, Claire M. Smith, Mariana O. Diniz, Bryan Williams, Paul A. Dalby, Gareth R. Williams, John Ward, Sudaxshina Murdan, Alejandro Arenas-Pinto, Jose Alejandro Guevara-Patiño, Delmiro Fernandez-Reyes

## Abstract

This work was conducted during the COVID-19 pandemic prior to the licensing of any vaccines against COVID-19. Although several COVID-19 vaccines are now commercially available, this research on the development of a combination Severe Acute Respiratory Syndrome-associated Coronavirus (SARS-CoV), SARS-coronavirus 2 (SARS-CoV-2), and Middle Eastern Respiratory Syndrome (MERS) is still relevant and shows how a combination vaccine can be designed, produced and rapidly tested in the laboratory. We present the development of a combination vaccine designed to provide immunity against Severe Acute Respiratory Syndrome-associated Coronavirus (SARS-CoV), SARS-coronavirus 2 (SARS-CoV-2), and Middle Eastern Respiratory Syndrome (MERS). The primary objective of this vaccine design is twofold: to mitigate the burden of coronavirus and to address the specific vulnerability of regions prone to recurrent MERS outbreaks. Our combination vaccine incorporates antigenic components from zoonotic sources, specifically SARS-CoV, SARS-CoV-2, and MERS. We assess the impact of combining different Spike protein’s S1 subunit antigens, due to its recognised immunogenic potential, and adjuvants on serum antibody titres, virus neutralizing capabilities, and inter-antigen immune responses. We report a robust and broad antibody response against SARS-CoV-2 and related coronaviruses, which was amplified by different adjuvant formulations, including alum, MPLA, CpG, and Squalene-in-Water Emulsion.

## INTRODUCTION

This work was conducted during the COVID-19 pandemic prior to the licensing of any vaccines against COVID-19. Although several COVID-19 vaccines are now commercially available, this research on the development of a combination Severe Acute Respiratory Syndrome-associated Coronavirus (SARS-CoV), SARS-coronavirus 2 (SARS-CoV-2), and Middle Eastern Respiratory Syndrome (MERS) is still relevant and shows how a combination vaccine can be designed, produced and rapidly tested in the laboratory.

Three zoonotic beta coronaviruses have been responsible in recent years, for human outbreaks of respiratory disease, including COVID-19. Developing vaccine strategies that can provide wider coverage against a broad range of coronaviruses and their variants are crucial to effectively prevent future coronaviruses outbreaks^1–5^.

Combination vaccines, like Measles, Mumps, and Rubella (MMR) and Diphtheria, Tetanus, Pertussis (DTaP), have many advantages, such as reduction in the numbers of vaccine administration events and visits to the vaccination centre, which can lead to higher vaccine coverage. In addition, combination vaccines offer reduced overall costs due to, for example, shared packaging, shipping, and administration costs. In this paper, we describe the development of a Severe Acute Respiratory Syndrome (SARS) -associated coronavirus (SARS-CoV), SARS-coronavirus 2 (SARS-CoV-2) and Middle Eastern Respiratory Syndrome (MERS) combination vaccine. Such a vaccine would be globally relevant and especially applicable to certain regions in the world, such as the Middle East region (e.g., Saudi Arabia)^6^, where repeated MERS outbreaks have occurred^7^. Episodic appearance of patients with MERS is via virus transmission from dromedary camels to camel workers (who develop a mild disease) to susceptible individuals who have not been exposed to camels, but who go on to develop significant disease ^8–11^. Thus, the latest MERS fatality in Saudi Arabia (as of 6 October) had reported no contact with camels^6^.

Given the recent global impact of SARS-CoV-2 and the relatively fewer variants of MERS-CoV, our research first delves into exploring and understanding the immunogenicity of various SARS-CoV and SARS-CoV-2 antigens. Building upon these findings, we then extend our study to incorporate MERS-CoV antigen, aiming to develop a combination vaccine that could mitigate the burdens of both diseases considering not only effectiveness of adjuvant formulations but more importantly their manufacturing scalability and availability.

Several antigens, related to the SARS-CoV-2 Wuhan and Alpha, Beta and Gamma variants (of concern at the time of the experiments), were first explored to address the mutating virus and emerging variants, which could reduce vaccine efficacy. Our second vaccine component addresses the zoonotic element, which is critical in preventing future epidemics and pandemics. The zoonotic component was represented by sequences related to SARS-CoV encountered in bats^12–14^. Our third vaccine component was the MERS antigen. For all three vaccine antigen components, the virus Spike protein, which enables viral attachment, fusion, and entry into host cells^15–18^, was the selected antigen. Specifically, the S1 subunit, containing the Receptor-Binding Domain (RBD) which is responsible for binding to the human receptor and initiating viral entry into human cells, was selected^15,16,19^. The RBD is highly immunogenic and elicits a strong neutralising antibody response, which can prevent viral entry and replication. Given that protein antigens are usually poorly immunogenic on their own, the fourth component of our vaccine was the adjuvant.

For the formulations studied, we show the antigen-specific and cross-reactive response to a wide panel of relevant antigens, as well as the virus-neutralizing titres to SARS-CoV-2. Our final vaccine, the SARS and MERS combination formulation, uses an industrially scalable squalene-based adjuvant.

## MATERIALS and METHODS

### Vaccine Formulation Antigens

The following eukaryotic expressed recombinant proteins were used as antigens for immunisation (see formulations Table 1). Recombinant SARS-CoV-2 Spike S1 subunit (Table 1, Wuhan^&^), SARS-CoV Spike S1 subunit (Table 1, Bat^&^) and SARS CoV2 S1 (Table 1, Alpha^@^, Beta^@^ and Gamma^@^ variants) were obtained from the Crick Institute (UK). Briefly, constructs were codon-optimized for expression in human cells and cloned under the control of the cytomegalovirus promoter for production of the recombinant proteins carrying a C-terminal extension containing human rhinovirus 14 3C protease recognition site followed by a Twin Strep tag. The vectors used for the expression was pcDNA3 for the alpha, beta, and gamma variants and pQ-3C2xStrep for the Bat and Wuhan proteins^20^. The proteins were then produced by transient transfection of Expi293 (Thermo Fisher Scientific) cells with endotoxin-free preparations of the corresponding DNA constructs using ExpiFectamine293 (Thermo Fisher Scientific). The cells were maintained in shake flasks in FreeStyle293 (Thermo Fisher Scientific) medium at 37°C in humidified 5% CO2 atmosphere. Twin-Strep-tagged proteins were captured on Strep-Tactin XT (IBA Lifesciences) affinity resin. Following extensive washes in TBSE [150 mM NaCl, 1 mM EDTA, and 25 mM tris-HCl (pH 8.0)], the proteins were eluted in 1× BXT buffer (IBA Lifesciences). Recombinant MERS-CoV spike S1 subunit protein was obtained from Bio-techne Ltd (UK)^20^.

**Table 1.**
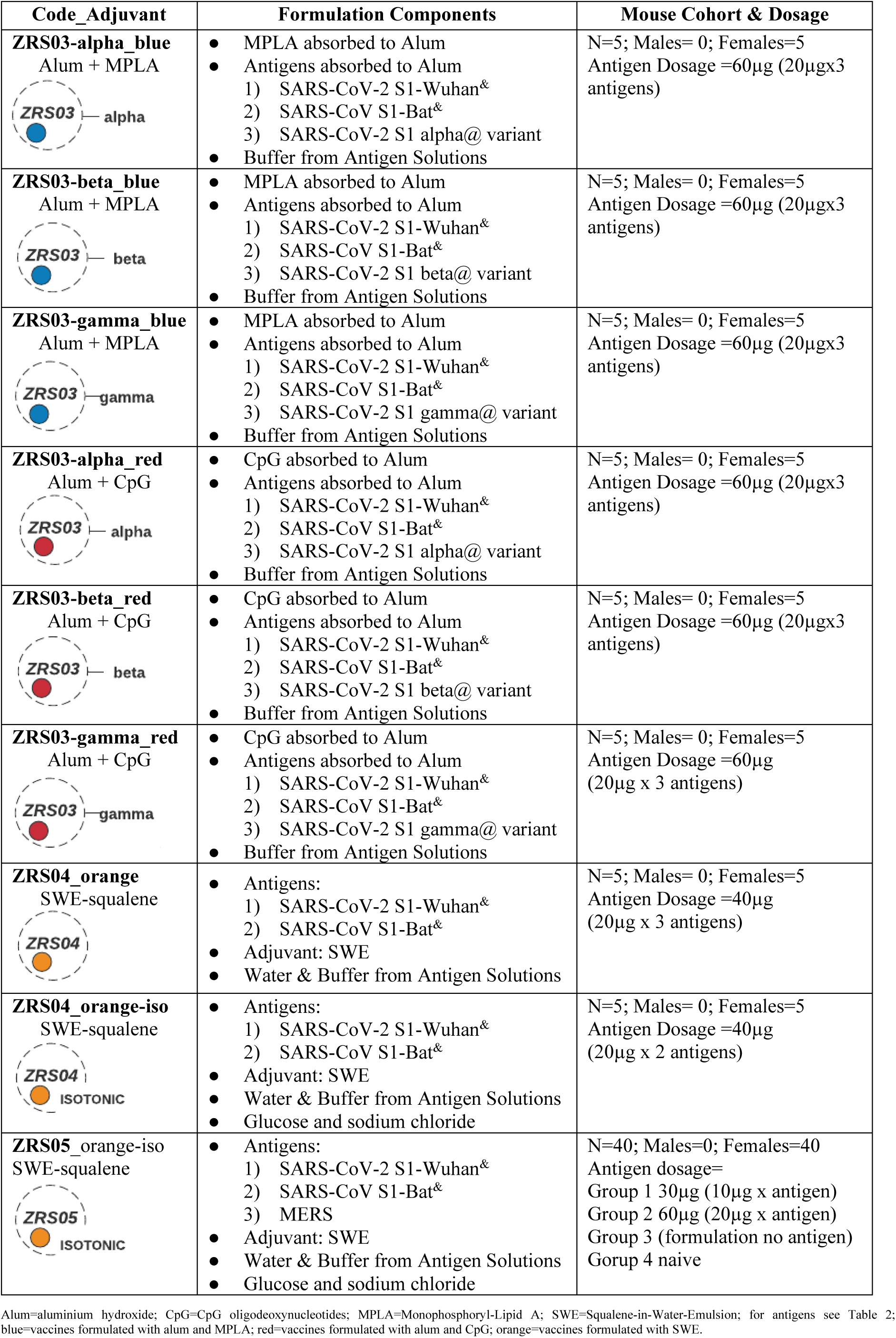
Vaccine formulations, adjuvants, dose, and characteristics of immunised cohorts.

| Code_Adjuvant | Formulation Components | Mouse Cohort & Dosage |
| --- | --- | --- |
| **ZRS03-alpha_blue**<br>Alum + MPLA | MPLA absorbed to Alum<br>Antigens absorbed to Alum<br>SARS-CoV-2 S1-Wuhan<sup>&</sup><br>SARS-CoV S1-Bat<sup>&</sup><br>SARS-CoV-2 S1 alpha@ variant<br>Buffer from Antigen Solutions | N=5; Males= 0; Females=5<br>Antigen Dosage =60µg (20µgx3 antigens) |
| **ZRS03-beta_blue**<br>Alum + MPLA | MPLA absorbed to Alum<br>Antigens absorbed to Alum<br>SARS-CoV-2 S1-Wuhan<sup>&</sup><br>SARS-CoV S1-Bat<sup>&</sup><br>SARS-CoV-2 S1 beta@ variant<br>Buffer from Antigen Solutions | N=5; Males= 0; Females=5<br>Antigen Dosage =60µg (20µgx3 antigens) |
| **ZRS03-gamma_blue**<br>Alum + MPLA | MPLA absorbed to Alum<br>Antigens absorbed to Alum<br>SARS-CoV-2 S1-Wuhan<sup>&</sup><br>SARS-CoV S1-Bat<sup>&</sup><br>SARS-CoV-2 S1 gamma@ variant<br>Buffer from Antigen Solutions | N=5; Males= 0; Females=5<br>Antigen Dosage =60µg (20µgx3 antigens) |
| **ZRS03-alpha_red**<br>Alum + CpG | CpG absorbed to Alum<br>Antigens absorbed to Alum<br>SARS-CoV-2 S1-Wuhan<sup>&</sup><br>SARS-CoV S1-Bat<sup>&</sup><br>SARS-CoV-2 S1 alpha@ variant<br>Buffer from Antigen Solutions | N=5; Males= 0; Females=5<br>Antigen Dosage =60µg (20µgx3 antigens) |
| **ZRS03-beta_red**<br>Alum + CpG | CpG absorbed to Alum<br>Antigens absorbed to Alum<br>SARS-CoV-2 S1-Wuhan<sup>&</sup><br>SARS-CoV S1-Bat<sup>&</sup><br>SARS-CoV-2 S1 beta@ variant<br>Buffer from Antigen Solutions | N=5; Males= 0; Females=5<br>Antigen Dosage =60µg (20µgx3 antigens) |
| **ZRS03-gamma_red**<br>Alum + CpG | CpG absorbed to Alum<br>Antigens absorbed to Alum<br>SARS-CoV-2 S1-Wuhan<sup>&</sup><br>SARS-CoV S1-Bat<sup>&</sup><br>SARS-CoV-2 S1 gamma@ variant<br>Buffer from Antigen Solutions | N=5; Males= 0; Females=5<br>Antigen Dosage =60µg (20µg x 3 antigens) |
| **ZRS04_orange**<br>SWE-squalene | Antigens:<br>SARS-CoV-2 S1-Wuhan<sup>&</sup><br>SARS-CoV S1-Bat<sup>&</sup><br>Adjuvant: SWE<br>Water & Buffer from Antigen Solutions | N=5; Males= 0; Females=5<br>Antigen Dosage =40µg (20µg x 3 antigens) |
| **ZRS04_orange-iso**<br>SWE-squalene | Antigens:<br>SARS-CoV-2 S1-Wuhan<sup>&</sup><br>SARS-CoV S1-Bat<sup>&</sup><br>Adjuvant: SWE<br>Water & Buffer from Antigen Solutions<br>Glucose and sodium chloride | N=5; Males= 0; Females=5<br>Antigen Dosage =40µg (20µg x 2 antigens) |
| **ZRS05_orange-iso**<br>SWE-squalene | Antigens:<br>SARS-CoV-2 S1-Wuhan<sup>&</sup><br>SARS-CoV S1-Bat<sup>&</sup><br>MERS<br>Adjuvant: SWE<br>Water & Buffer from Antigen Solutions<br>Glucose and sodium chloride | N=40; Males=0; Females=40<br>Antigen dosage=<br>Group 1 30µg (10µg x antigen)<br>Group 2 60µg (20µg x antigen)<br>Group 3 (formulation no antigen)<br>Group 4 naive |

### Vaccine Formulation Adjuvants

The following adjuvants were used in our vaccine formulations (see formulations Table 1). Vaccigel alum adjuvant (2% aluminium hydroxide) was purchased from 2BScientific (UK). CpG oligodeoxynucleotides mouse (1018) adjuvant was obtained from Bio-techne Ltd (UK). MPLA (PHAD) was purchased from Sigma-Aldrich (Switzerland). Squalene* oil-in-water emulsion adjuvant SEPIVAC Squalene-in-Water Emulsion (SWE)™ was obtained from SEPPIC (UK). Unless otherwise stated, all reagents and chemicals used in this study were of analytical grade.

### Vaccine Formulations of ZRS03

The protein antigens used for immunization in ZRS03 cohort (Table 1, Wuhan^&^, Bat^&^, Alpha^@^, Beta^@^ and Gamma^@^) and the adjuvants MPLA & CpG were adsorbed to alum before the immunizations. Adsorption to alum, for each protein and adjuvant, was conducted separately to avoid adsorption competition among the adsorbates. Specific volumes of each of the protein, CpG or MPLA solutions was placed in a glass vial to which alum was added. The mixtures were allowed to mix on a rotator for 2 hrs at 4 ͦ C, after which, the protein-alum mixtures were transferred to the MPLA-alum or CpG-alum mixtures. These formulations were left overnight at room temperature prior to use the next day.

### Vaccine Formulations of ZRS04

Two vaccine formulations (Table 1), one adjusted for isotonicity and one hypertonic (adding the squalene-based adjuvant make the vaccine mix hypertonic) were prepared to determine the importance of formulation isotonicity. Antigen solutions (Table 1, Wuhan^&^ and Bat^&^ proteins), water OR an aqueous solution containing glucose (10% w/v) and sodium chloride (0.9% w/v) and finally SWE adjuvant was placed into a glass vial. The vial was then gently inverted 10 times to mix the components following SEPIVAC Squalene-in-Water Emulsion (SWE)™ manufacturer instructions SEPPIC (UK), and then kept in the fridge until used the next day.

### Vaccine Formulation of ZRS05

An isotonic vaccine formulation was prepared by mixing antigen solutions (Table 1, Wuhan^&^, Bat^&^ and MERS proteins), an aqueous solution containing glucose (10% w/v) and sodium chloride (0.9% w/v) and finally SWE adjuvant, in a glass vial (Table 1). The vial was then gently inverted 10 times to mix the components following SEPIVAC Squalene-in-Water Emulsion (SWE)™ manufacturer instructions SEPPIC (UK), and then kept in the fridge until used the next day.

### Mice Cohorts and Immunisation Protocols

Female Balb/C mice, 20g in weight (8-15 weeks old) were purchased from Charles River Lab (UK). Following acclimatisation for a week, mice in groups of 5 (ZRS03, ZRS04) or 10 (ZRS05) were immunized subcutaneously with 100 microlitre of the vaccine formulation as shown in Table 1. Each mouse received prime and booster vaccine doses, and bled at intervals, as shown in the vaccination scheme (Figure 1), to collect serum for measurement of antibody and virus neutralization titres.

**Figure 1.**
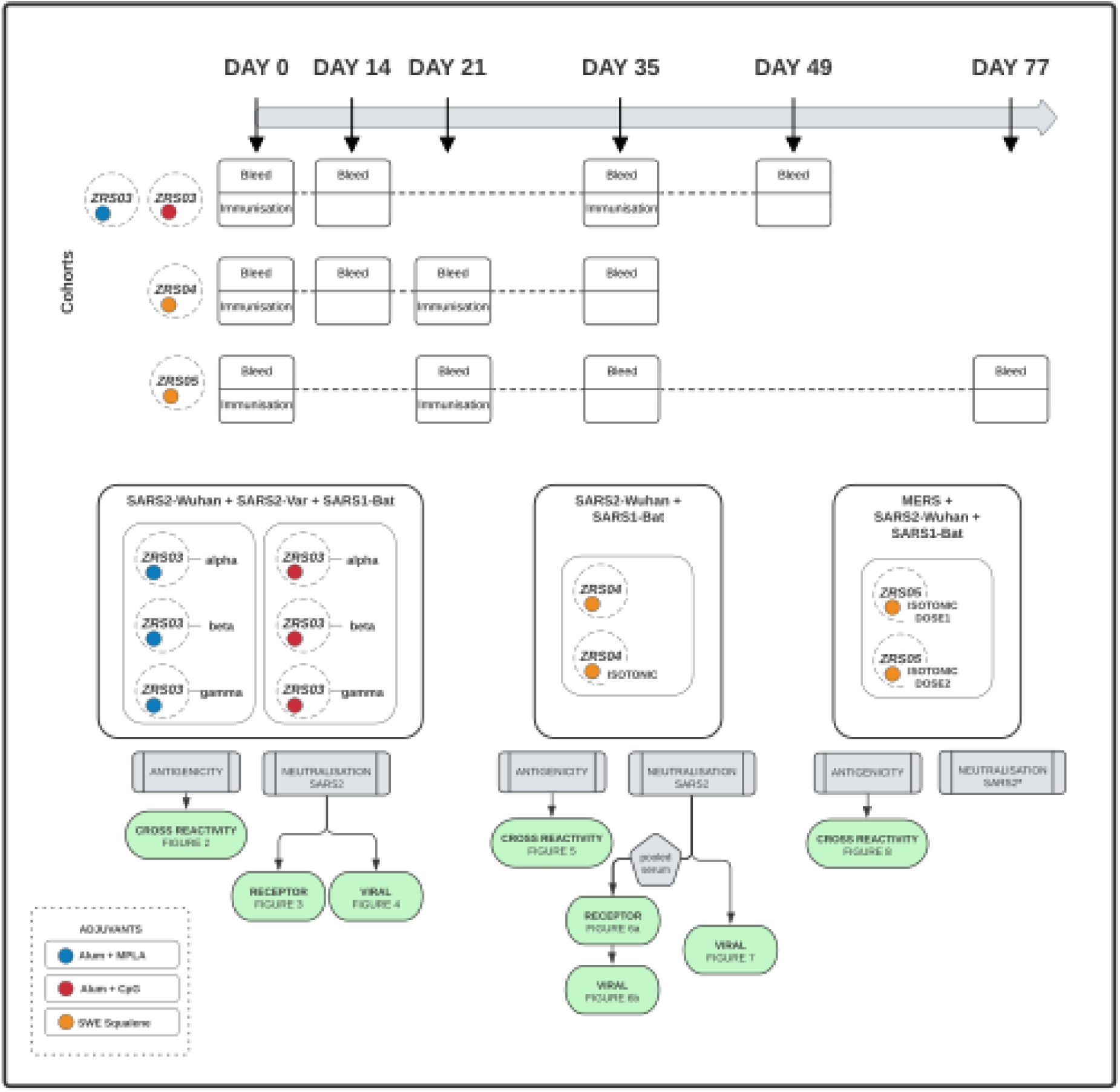
Mice immunisation and bleed protocol, vaccine antigens, adjuvant, and formulation overview. ZRS=Zoogenic Recombinant SARS; blue=vaccines formulated with alum and MPLA; red=vaccines formulated with alum and CpG; orange=vaccines formulated with SWE. ZRS03=Zoogenic Recombinant SARS version 3 containing as antigens the S1 sub-unit of (SARS2-Wuhan + SARS2-variant [either alpha, beta or gamma] + SARS1-bat) (ZRS04, ZRS05 see Table 1). Alum=aluminium hydroxide; CpG=CpG oligodeoxynucleotides; MPLA=Monophosphoryl-Lipid A; SWE=Squalene-in-Water-Emulsion.

Briefly, for ZRS03 cohort, mice were immunised at day 0 and day 35 for ZRS03, except for ZRS03-beta group, which did not receive a booster. Blood (100µl) were collected at days 0, 14, and 35 and mice culled for a terminal bleed at day 42 (Table 1, Figure 1). Cohort ZRS04 was immunised at days 0 and 21, and 100µl of blood were collected at day 0,14, and day 35 terminal bleed (Table 1, Figure 1). Mice from cohort ZRS05 were immunised at day 0 and 21 and 100µl of blood were collected at days 0,21,35 and 77 terminal bleed (Table 1, Figure 1).

### Serum IgG ELISAs

In-house indirect enzyme-linked immunosorbent assay (ELISAs) were developed for the determination of IgG antibodies to the target antigens. The 96-well ELISA plates (Nunc Maxisorp) were coated with 100 µL per well of antigen solution at 1 µg/ml in coating buffer (pH 9.4, Thermo BupH™ Carbonate-Bicarbonate Buffer), incubated overnight at 4 °C for adsorption. A standard curve of mouse IgG isotype control was also adsorbed (Thermo, 02-6502). The assay buffer (1x eBioscience™ ELISA/ELISPOT Diluent, 00-4202-56) was used for blocking (1 hour at RT) and as sample diluent (1:500 dilution of serum and controls). A SARS2 spike primary mouse antibody (CE5, Native Antigen Company, MAB12443) was included as a positive control, and normal mouse serum (Invitrogen, 10410) used as a negative control. Samples were added to the plate in triplicate wells of 100 µL and incubated for 2 hours at RT. All wells were washed with 200 µL of wash buffer (0.05% PBS-T) for three 5-minute incubations on a shaking platform. A 2-hour incubation at RT with 100 µL of horseradish peroxidase (HRP)-conjugated goat anti-mouse IgG antibody (1:2000 in sample diluent, Merck A4416-1M) was used for detection, followed by washing as before. Wells were developed by addition of 100 µL of TMB substrate (Thermo N301) for 10 minutes, followed by 100 µL of stop solution (0.16M sulfuric acid, Thermo N600). A FLUOstar Optima microplate reader (BMG Labtech) was used to measure optical density (OD) at an absorbance of 450nm.

The antigens used for determination of the resulting post-immunisation antibody response are described in Table 2.

**Table 2.** Antibodies in experimental and control cohorts were tested for the ability to recognise.

| **Virus** | **Variant Code** | **Subunit** | **Manufacturer Code** |
| --- | --- | --- | --- |
| SARS-CoV-2 | Wuhan<sup>&</sup> | Spike S1 subunit<br>(Used in formulation) | [21] |
| SARS-CoV | Bat<sup>&</sup> | Spike S1 subunit<br>(Used in formulation) | [21] |
| SARS-CoV | SARS 1<sup>@</sup> | Spike S1 subunit<br>(Used in formulation) | Bio-Techne(R&D)<br>10783-cv |
| SARS-CoV-2 | Wuhan<sup>@</sup> | Spike S1 subunit | Bio-Techne(R&D)<br>10569-cv |
| SARS-CoV-2 | Bat<sup>@</sup> | Spike S1 subunit RBD | Bio-Techne(R&D)<br>10593-cv |
| SARS-CoV-2 | Alpha<sup>@</sup><br>(UK) B.1.1.7 | Spike | Bio-Techne(R&D)<br>10748-cv |
| SARS-CoV-2 | Beta<sup>@</sup><br>(S. Africa) B.1.351 | Spike | Bio-Techne(R&D)<br>10777-cv |
| SARS-CoV-2 | Gamma<sup>@</sup><br>(Brazil) | Spike | Bio-Techne(R&D)<br>10795-cv |
| SARS-CoV-2 | Delta<sup>@</sup><br>(India) B.1.617.2 | Spike | Bio-Techne(R&D)<br>10942-cv |
| SARS-CoV-2 | Kappa<sup>@</sup><br>(India) | Spike | Bio-Techne(R&D)<br>10861-cv |
| SARS-CoV-2 | Omicron<sup>@</sup><br>B.1.1529 | Spike | Stratech (Sino-Bio)<br>40591-V08H4 |
| MERS | MERS | Spike | Bio-Techne(R&D)<br>10739-cv |

### Antibody Neutralisation Assay

We used a modified protocol developed from an anti-SARS-CoV-2 Neutralising Antibody Titre Serologic Assay Kit (ACRO Biosystems, TAS-K022). In brief, 55 µL of diluted serum sample was mixed with 55 µL of HRP-conjugated SARS-CoV-2 Spike RBD (0.3 ug/mL) and incubated for 30 minutes. These mixtures were immediately transferred to wells pre-coated with human ACE2 and incubated 1 hour at 37 °C. The wells were washed four times with a wash buffer (from ACRO Biosystems kit), each for 1 minute on a shaking platform. Wells were developed by addition of 100 µL of substrate solution (from ACRO Biosystems kit) incubated at 37 °C for 20 minutes, before 50 µL of stop solution was added. The optical density of each well was measured at 450nm using a FLUOstar Optima microplate reader (BMG Labtech).

### Viral Neutralisation Assay

Vero cells were plated the day before the experiment at 8,000 cells/well in black 96 well plates. On the day of the experiment, serum samples were diluted 3-fold (8 dilutions) in triplicates, starting from 1:30. Virus (SARS-CoV2/England/2/2020; BEI Resources) was added at an MOI of 0.5, and virus plus serum dilutions were incubated for 1 hour at 37 °C. At the end of the incubation, the virus + serum mixtures were added to the assay cells and incubated for 6 hours at 37°C, 5% CO2, after which cells were fixed and stained for a viral antigen using a rabbit anti-N antibody. Images were acquired using a high-content Cell Insight instrument and the percentage of infected cells was calculated over the total number of cells. Percentages of inhibition are normalized to the plate internal controls (uninfected untreated and uninfected untreated cells). WHO positive and negative sera were included as assay controls.

### Assessment of T-Cell responses

To access T-cell responses, ELISPOT was carried on cells from spleens obtained seven days post-boost terminal-bleed of mice. ELISPOT was carried out using ELISpot Plus: Mouse IFN-γ (HRP) (MABTECH) and following manufacturer’s instructions. The following MHC-II peptides, termed CD4 peptides pools were used: CEFTA peptide pool human CD4 T-cell (MABTECH); PepMix MERS-CoV 336 peptides (jpt, PM-MERS-CoV-S-1); SARS-CoV-2 S1 scanning pool 166 peptides (MABTECH) (Supplementary Figure 1). MHC-I peptides termed CD8 peptide pools were designed and synthetised (Supplementary Figure 1).

## RESULTS

Here we report the effect of the inclusion of the different relevant antigens and adjuvants on a) antigen-specific and cross-reactive serum antibody levels; b) virus neutralising antibody titres and c) any positive, neutral, or negative impact of one antigen on the immune response of another antigen. The overall vaccination strategy is described in Figure 1 and Table 1.

### Zoonotic Coronaviruses Multivalent Immunisation Strategy

Three vaccines named Zoonotic Recombinant SARS-CoV (ZRS), ZRS03, ZRS04 and ZRS05 were explored to study the widening of the elicited antibody and neutralising immune response (Figure 1 and Table 1).

First, our initial formulation ZRS03 includes three eukaryotic-system expressed recombinant Spike S1 subunits: SARS-CoV-2 Wuhan^21^, one of the three SARS-CoV-2 variants of concern (alpha^21^, beta^21^, gamma^21^) and a zoonotic component of SARS-CoV present in bats^21^. The resulting combinations ZRS03-alpha, ZRS03-beta and ZRS-gamma were formulated either with alum-MPLA (Figure 1 and Table 1, -blue) or alum-CpG (Figure 1 and Table 1, -red) adjuvants. As expected, we did not observe any severe adverse effect in the experimental animal cohorts.

To assess the response range to emerging SARS-CoV-2 variants a vaccine incorporating only two antigens, SARS-CoV-2 Wuhan and SARS-CoV Bat, was then designed and formulated with the squalene-based adjuvant SWE^22–25^ ZRS04 (Figure 1 and Table 1 -orange) which, besides being previously used in experimental and clinical setting, is free of patent rights, scalable, and cost-effective for large population vaccination. ZRS04 formulation was developed in both isotonic and non-isotonic versions (Figure 1 and Table 1 -orange). We did not observe any severe adverse effect in the experimental animal cohorts when using this adjuvant.

Finally, we turned our attention to incorporate a third antigen to provide a rationale for a vaccine that covers SARS-CoV, SARS-CoV-2 and MERS with the justification that sequence diversity given using this combination would provide a wider response that could contribute to prevent the jump of the zoonotic MERS into humans. Given our results with the squalene-based adjuvant, we formulated the ZRS05 vaccine using SWE in isotonic conditions and exploring two dosage schemes (Figure 1 and Table 1 -orange). We did not observe any severe adverse effect in the experimental animal cohorts when using this adjuvant.

### Immunisation with ZRS03 vaccine formulations adjuvanted with either alum-MPLA or alum-CpG elicits pan-variant neutralising antibody immune response to SARS-CoV-2

First, we investigated whether the ZRS03-alpha, ZRS03-beta and ZRS03-gamma combinations induced a pan-variant neutralising response against SARS-CoV-2 and related coronaviruses. We found that two-weeks post-boost serum of animals vaccinated with ZRS03 (Figure 1) formulated either with alum-MPLA (Figure 2, -blue) or alum-CpG (Figure 2, -red) adjuvants elicited robust (>2 log over base level) antibody responses against all SARS-CoV antigens tested. No cross-reactivity was detected against the MERS antigen with either adjuvant. Alum-MPLA (Figure 2, -blue) and alum-CpG (Figure, 2-red) adjuvant combinations were equally effective, with no difference in serum antibody levels. A difference in the two adjuvant combinations was, however, found when *in vitro* cell-free antigen binding S1-RBD to hACE2 receptor was investigated (Figure 3). At two weeks post-boost, we observed a 50% neutralisation of binding in serum dilutions of 1:810 for both ZRS03-alpha and ZRS03-beta and 1:90 for ZRS03-gamma when immunisation involved the alum-MPLA adjuvant (Figure 3a, -blue). However, with the use of the alum-CpG adjuvant, serum dilutions required were 1:90 for ZRS03-alpha, and 1:270 for both ZRS03-beta and ZRS03-gamma (Figure 3b, -red).

**Figure 2.**
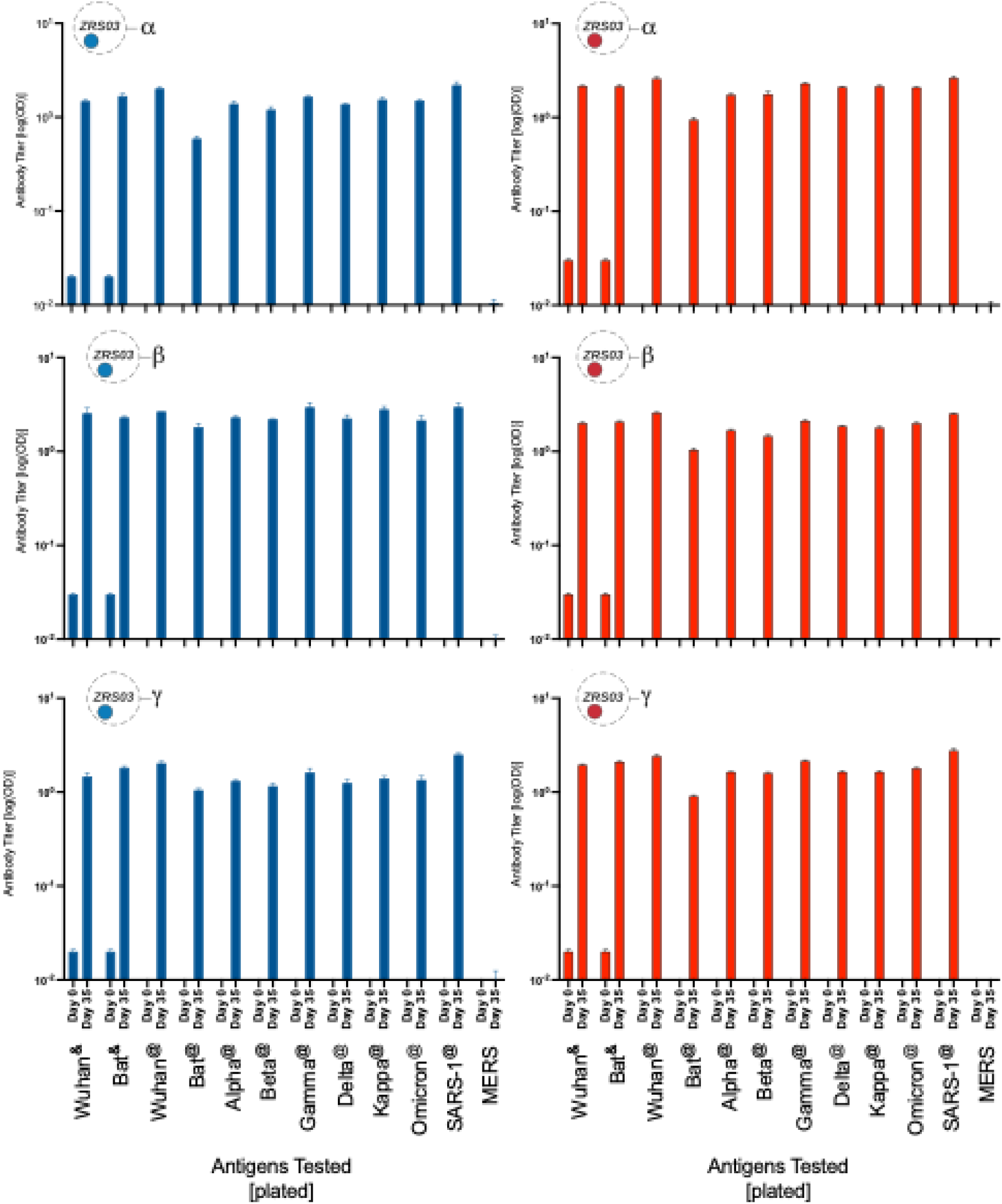
Serum antibody immunogenicity of ZRS03. ZRS=Zoogenic Recombinant SARS; blue=vaccines formulated with alum and MPLA; red=vaccines formulated with alum and CpG; ZRS03=Zoogenic Recombinant SARS version 3 containing as antigens the S1 sub-unit of (SARS2-Wuhan + SARS2-variant [either alpha, beta or gamma] + SARS1-bat) (see Table 1). Antigens tested see (Table 2). Alum=aluminium hydroxide; CpG=CpG oligodeoxynucleotides; MPLA=Monophosphoryl-Lipid A.

**Figure 3.**
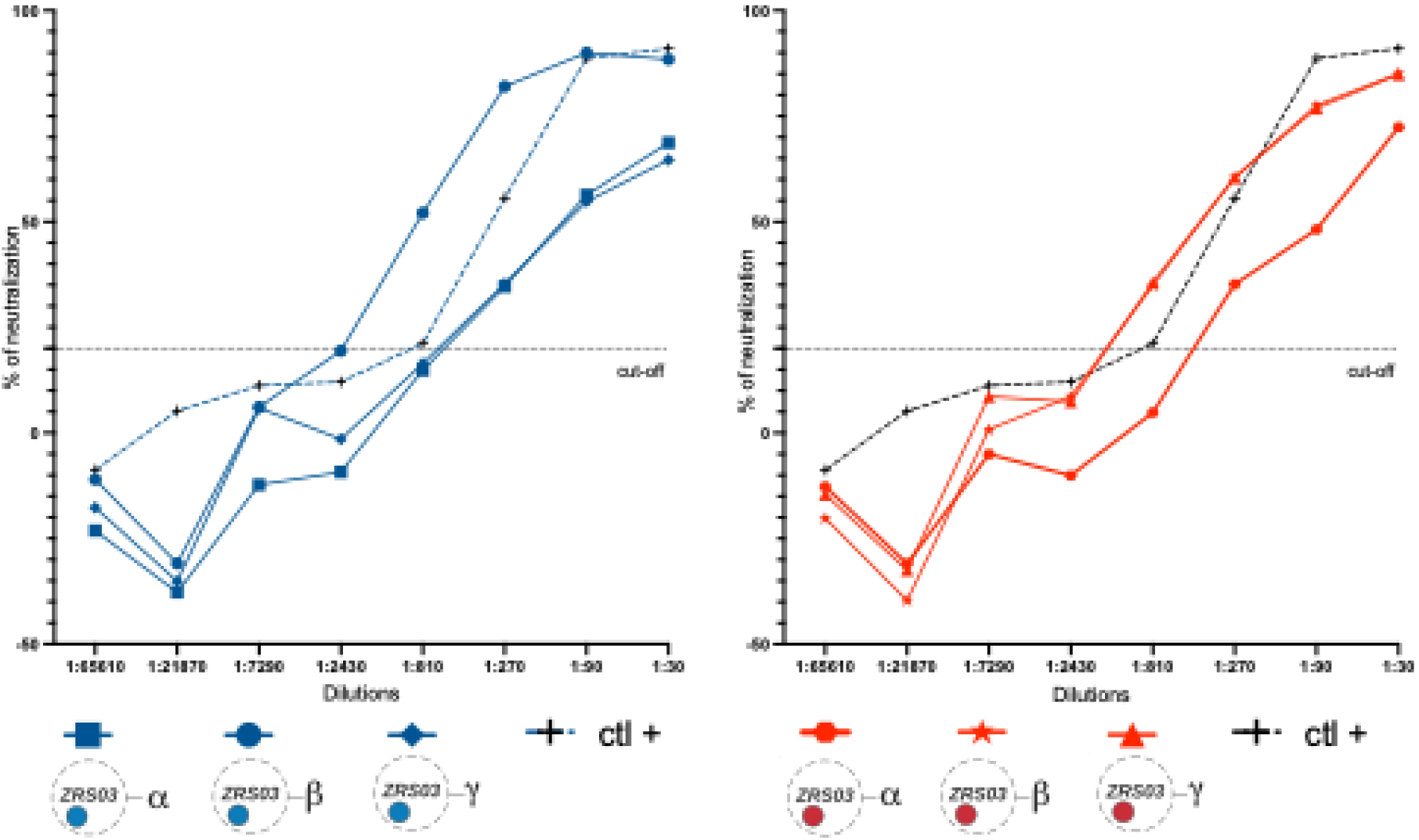
Neutralisation of ACE2 receptor SARS-CoV2 interaction by serum of ZRS03 immunised mice. ZRS=Zoogenic Recombinant SARS; blue=vaccines formulated with alum and MPLA; red=vaccines formulated with alum and CpG; ZRS03=Zoogenic Recombinant SARS version 3 containing as antigens the S1 sub-unit of (SARS2-Wuhan + SARS2-variant [either alpha, beta or gamma] + SARS1-bat) (see Table 1). Alum=aluminium hydroxide; CpG=CpG oligodeoxynucleotides; MPLA=Monophosphoryl-Lipid A.

All ZRS03 achieved SARS-CoV-2 viral neutralising titres (Figure 4, Table 3). Viral neutralisation assays within two-weeks post-boost serum from ZRS03-alpha and ZRS-gamma immunised mice elicited a 50% Tissue Culture Infectious Dose (TCID50) of 1:1620 and ZRS-gamma elicited lower 1:540 for when the alum-MPLA was used (Figure 4a, -blue; Table 3). Interestedly, the alum-CPA resulted in lower TCID50 for ZRS03-alpha (1:43740) and improved TCID50 for ZRS03-beta and ZRS03-gamma at 1:4860 and 1:1620 respectively (Figure 4b, -red; Table 3).

**Figure 4.**
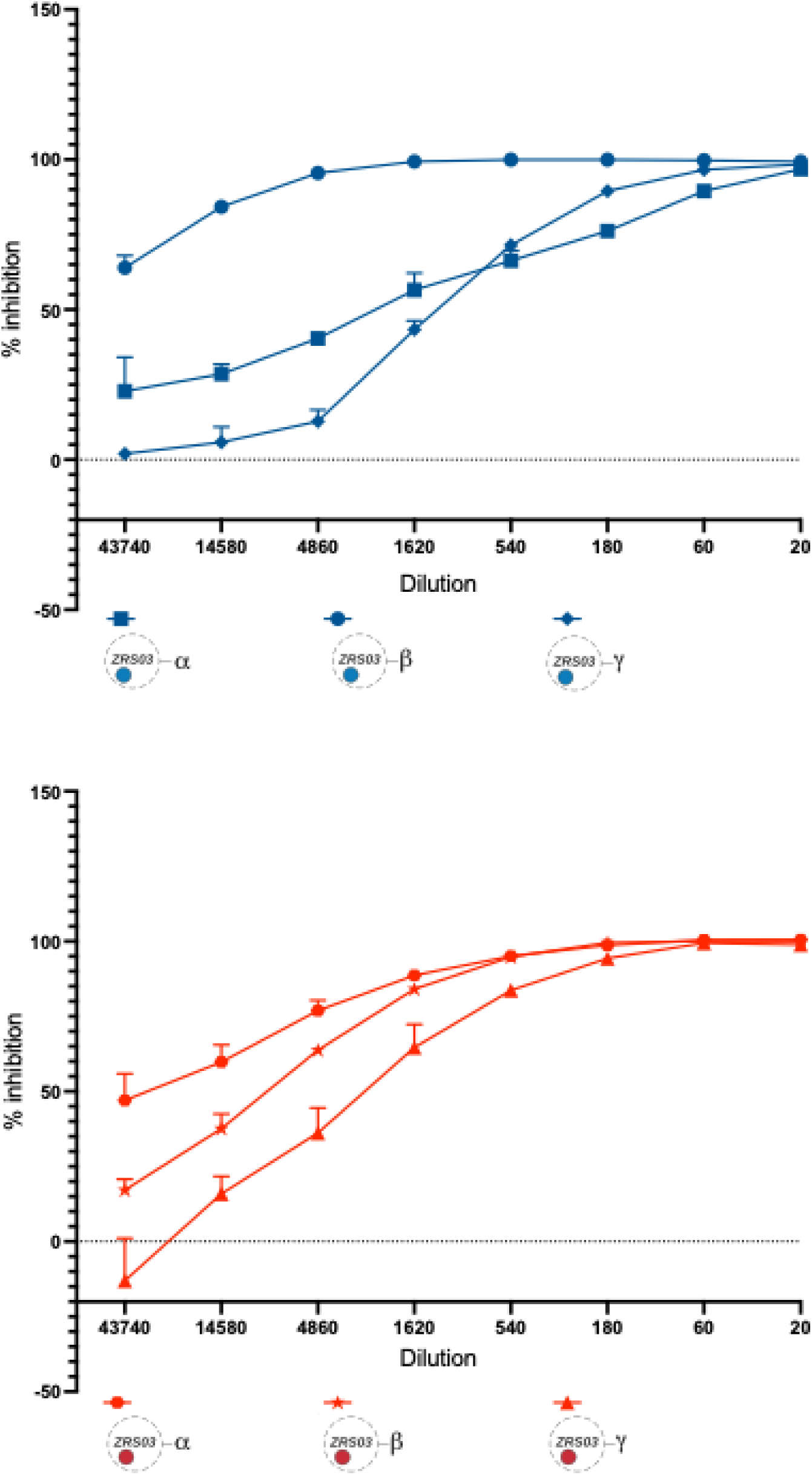
SARS-CoV2 *in vitro* viral neutralisation by serum of ZRS03 immunised mice. ZRS=Zoogenic Recombinant SARS; blue=vaccines formulated with alum and MPLA; MP; red=vaccines formulated with alum and CpG; ZRS03=Zoogenic Recombinant SARS version 3 containing as antigens the S1 sub-unit of (SARS2-Wuhan + SARS2-variant [either alpha, beta or gamma] + SARS1-bat). (see Table 1). Alum=aluminium hydroxide; CpG = CpG oligodeoxynucleotides; MPLA=Monophosphoryl-Lipid A.

**Table 3.** Neutralisation of studied vaccine formulations.

| Neutralisation (%)<br>System | Vaccine Formulation |  |  |  |  |  |
| --- | --- | --- | --- | --- | --- | --- |
|  | Adjuvant | ZRS03-alpha | ZRS03-beta | ZRS03-gamma | ZRS04<br>isotonic | ZRS04<br>hypotonic |
| **50%**<br>**Spike S1-hACE2**<br>**(post-boost)** | Alum-MPLA | 1:810 | 1:90 | 1:90 | - | - |
|  | Alum-CpG | 1:30 | 1:270 | 1:270 | - | - |
|  | SWE | - | - | - | 1:290 | 1:90 |
| **50%**<br>**Viral**<br>**(post-boost)** | Alum-MPLA | <1:43740 | 1:1620 | 1:540 | - | - |
|  | Alum-CpG | >1:43740 | 1:4860 | 1:1620 | - | - |
|  | SWE | - | - | - | 1:43740 | 1:43740 |
| **50%**<br>**Viral**<br>**(post-prime)** | Alum-MPLA | - | - | - | - | - |
|  | Alum-CpG | - | - | - | - | - |
|  | SWE | - | - | - | - | >1:60 |

### ZRS04 vaccine formulations adjuvanted with SWE squalene induces pan-variant immune response to SARS-CoV-2

Due to global difficulty with supplies and cost of MPLA and CpG adjuvants, a cost-effective open-access squalene based alternative adjuvant (Sepic SWE) was investigated. Mice were immunised with a combination of SARS-CoV-1 and SARS-CoV-2 Wuhan Spike S1 subunits formulated with SWE, either in an isotonic system (as recommended by the Sepic) or a hypotonic medium. One-week post-boost pooled serum from animals immunised with the squalene-based adjuvant system elicited a pan-variant antibody response against SARS-CoV-2, with a robust (>2 log) increase in antibody levels against all SARS antigens tested (Wuhan, Bat, Alpha, Beta, Gamma, Delta, Kappa, and Omicron), with no difference between the isotonic and hypotonic formulations (Figure 5a-b, -orange). Some differences were found in neutralisation of cell-free *in vitro* antigen binding S1-RBD to hACE2 receptor assays, with IC50 titres of 1:290 (ZRS04 isotonic) and 1:90 (ZRS04 hypotonic) (Figure 6a, -orange).

**Figure 5.**
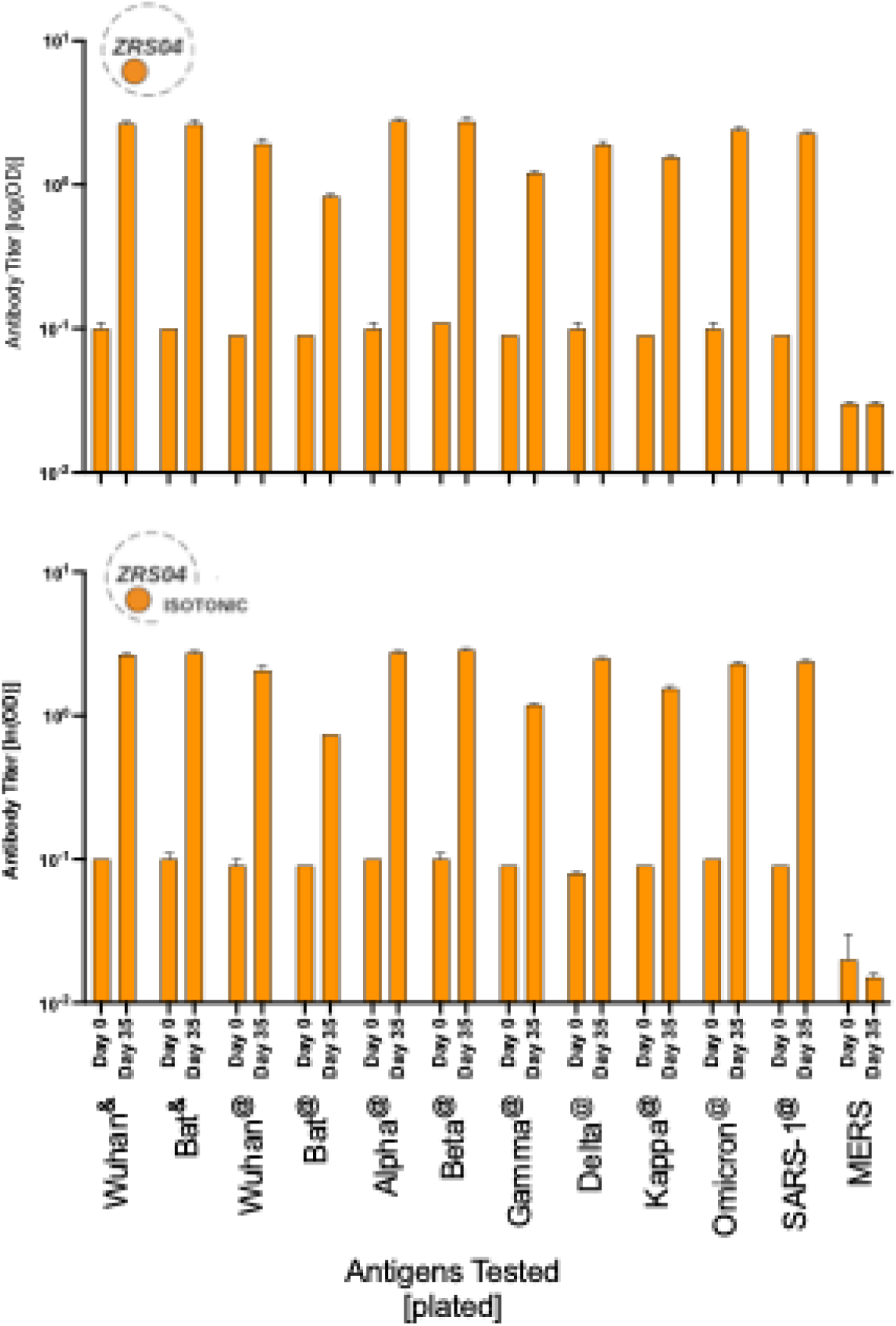
Serum antibody immunogenicity of ZRS04. ZRS=Zoogenic Recombinant SARS; orange=vaccines formulated with SWE; ZRS04=Zoogenic Recombinant SARS version 4 containing as antigens the S1 sub-unit of (SARS2-Wuhan + SARS1-bat) (see Table 1). Antigens tested see Table 2. SWE=Squalene-in-Water-Emulsion.

**Figure 6.**
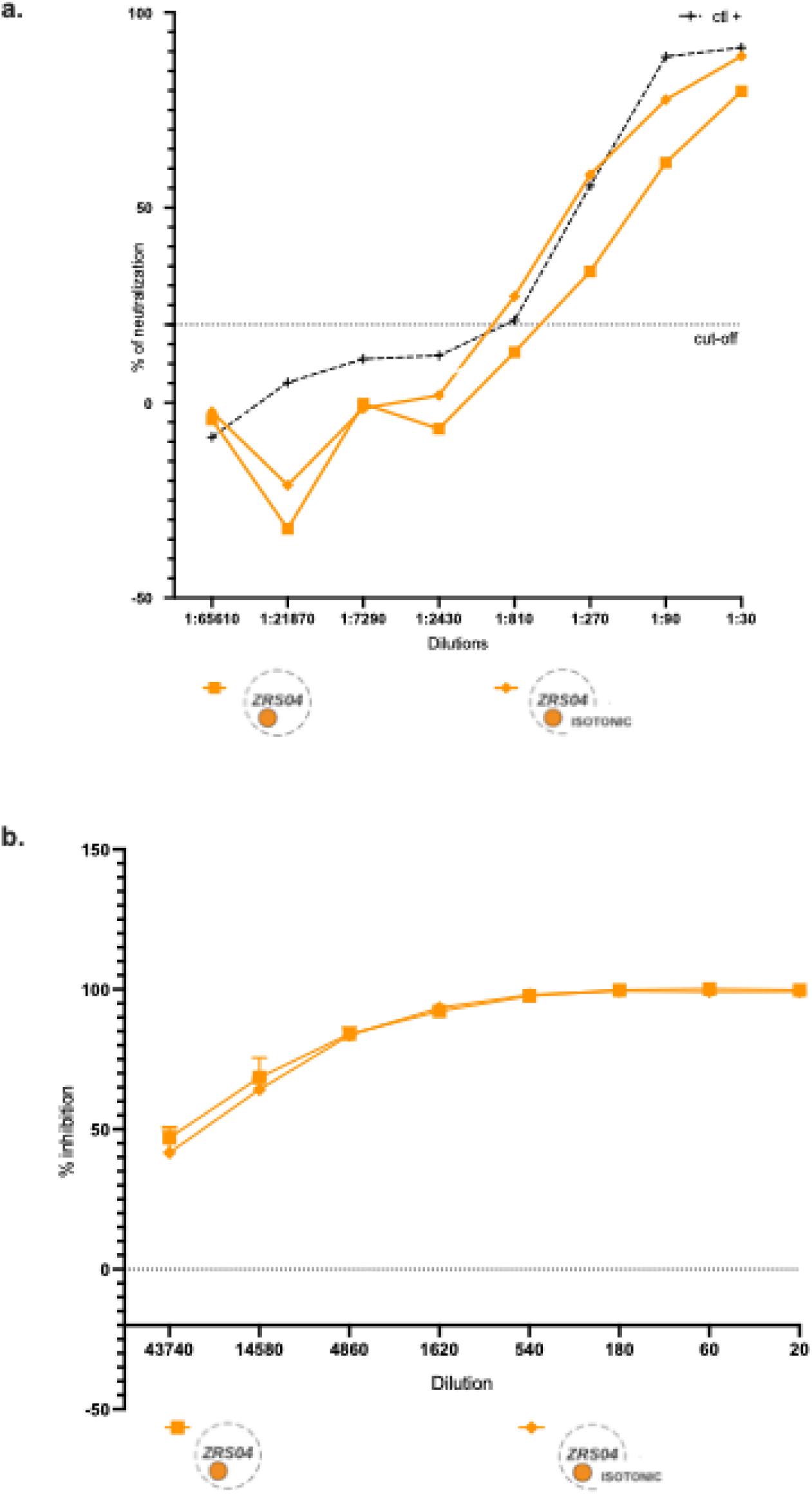
Neutralisation by pooled serum of ZRS04 immunised mice. **a.** Neutralisation of ACE2 receptor SARS-CoV2 interaction; **b.** SARS-CoV2 *in vitro* viral neutralisation. ZRS = Zoogenic Recombinant SARS; orange = vaccines formulated with SWE; ZRS04 = Zoogenic Recombinant SARS version 4 containing as antigens the S1 sub-unit of (SARS2-Wuhan + SARS1-bat) (see Table 1). SWE = Squalene-in-Water-Emulsion.

One-week post-boost pooled serum from animals immunised with both ZRS04 formulations showed high SARS-CoV-2 viral neutralisation, with 100% inhibition at titre of 1:540 and 50% inhibition at titre of 1:43740 (Figure 6b, -orange). Individual animal serum two-weeks post-prime serum from animals immunised with ZRS04 isotonic showed SARS-CoV-2 50% viral neutralisation at a titre of 1:180 but never achieved 100% neutralisation at highest titre of 1:20 (Figure 7). The terminal bleed serum of such animal cohort two-weeks post-boost achieved 50% and 100% viral neutralisation at titres of 1:43740 and 1:540 respectively (Figure 7) showing the adequate boosting of an efficient immune response.

**Figure 7.**
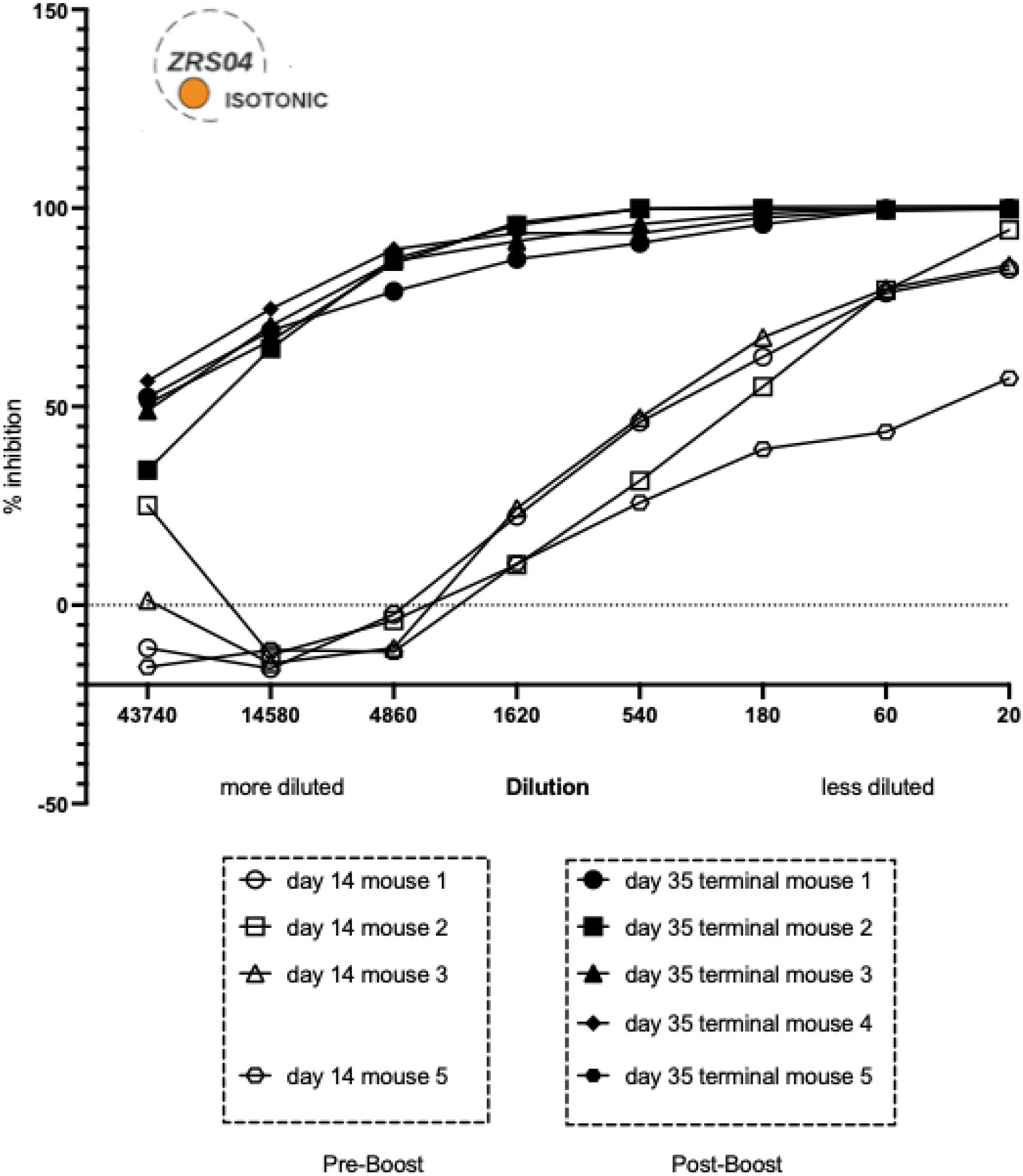
SARS-CoV2 *in vitro* viral neutralisation by serum of ZRS04-isotonic immunised mice pre- and post-boost. ZRS = Zoogenic Recombinant SARS; orange = vaccines formulated with SWE; ZRS04 = Zoogenic Recombinant SARS version 4 containing as antigens the S1 sub-unit of (SARS2-Wuhan + SARS1-bat) (see Table 1). SWE = Squalene-in-Water-Emulsion.

### Inclusion of MERS-CoV Spike S1 subunit in ZRS05 is essential for MERS-specific antibody responses and does not inhibit generation of pan-variant SARS-CoV-2-specific antibodies

Antibodies elicited by immunisation with SARS-CoV-1 and SARS-CoV-2 Spike S1 subunits in the presence of alum-MPLA or alum-CpG or SWE adjuvant do not induce cross-reactivity to the MERS-CoV Spike S1 antigen (Figures 2, 5). The Spike proteins of SARS and MERS coronaviruses are sufficiently different that a SARS vaccine is unlikely to protect against MERS. Therefore, we then explored ZRS05 a formulation based on ZRS04 SWE isotonic that includes MERS-CoV S1 subunit (Figure 1, Tables 1, 2). A total of 20 mice were vaccinated with ZRS05, with two different escalating dose regimens: 30µg (10µg of antigen, Group 1) and 60µg (20µg of antigen, Group 2). As controls, ten mice where vaccinated with antigen free formulation (Group 3) or left untreated (Group 4) (Table 1). Addition of a MERS-CoV Spike S1 subunit is necessary for the generation of MERS-specific antibodies, (Figure 8). Figure 8 also shows that the addition of a MERS antigen to SARS-CoV-1 and SARS-CoV-2 Spike S1 subunits in an isotonic SWE formulation did not have a negative impact on the generation of SARS-CoV-2-specific antibodies, and a pan-variant antibody response was found (Figure 8). This is promising for the development of a SARS-MERS combination vaccine relevant to middle east potential risk of MERS.

**Figure 8.**
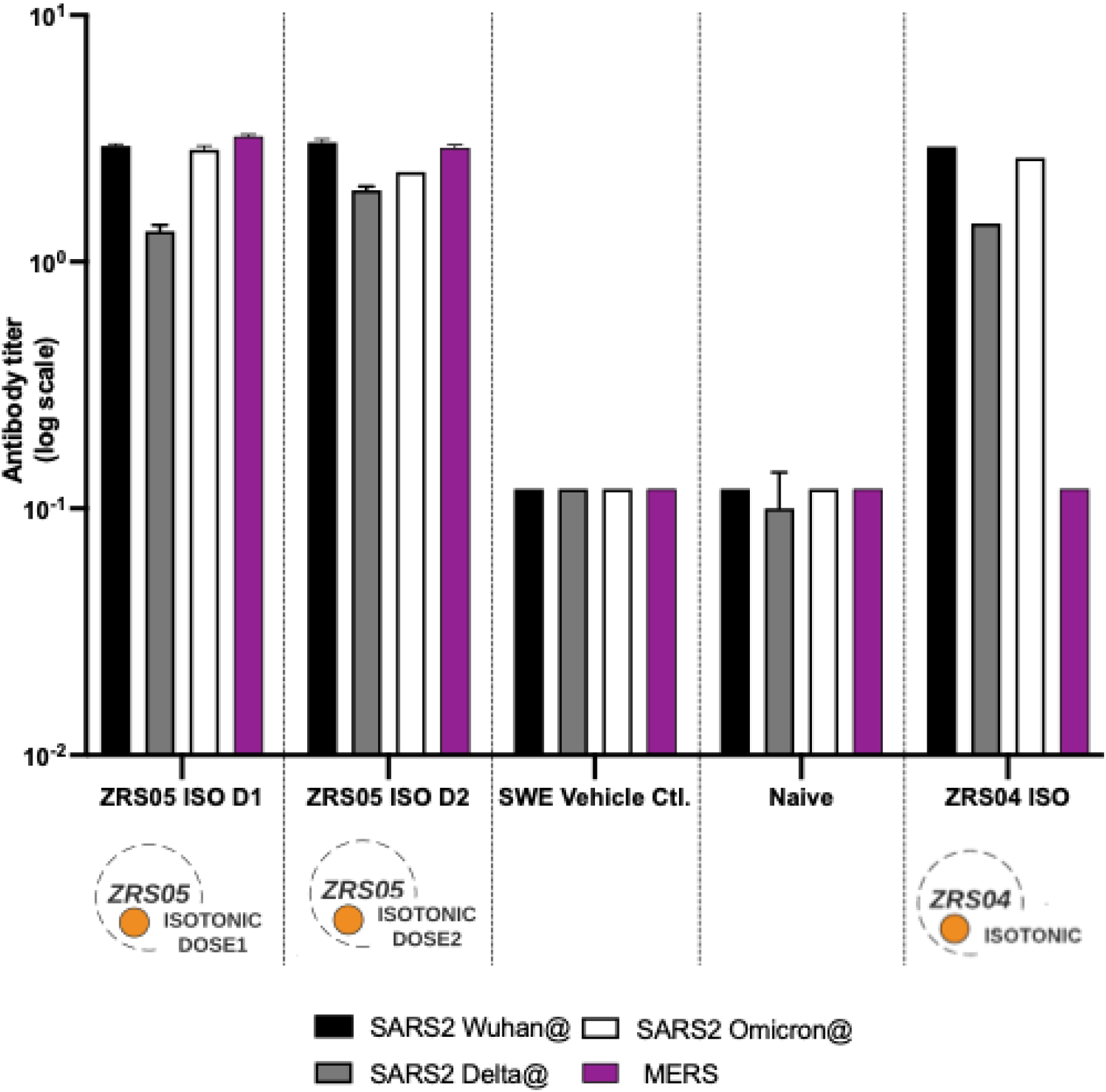
Serum antibody immunogenicity of isotonic ZRS05. ZRS=Zoogenic Recombinant SARS; orange=vaccines formulated with SWE; ZRS05=Zoogenic Recombinant SARS version 5 containing as antigens the S1 sub-unit of (SARS2-Wuhan + SARS1-bat) and MERS (see Table 1). ISO=isotonic formulation; Antigens tested see Table 2. SWE=Squalene-in-Water-Emulsion.

### ZRS05 SWE squalene-based formulation does not elicit a T-cell response

As ZRS05 was our selected combined formulation, we used IFN-γ Enzyme-Linked Immuno Spot (ELISpot) assay to interrogate whether squalene adjuvanted-ZRS05 immunisation induced CD4 and CD8 T cell responses to SARS-CoV-2 and MERS coronavirus (MERS-CoV)-derived epitopes. A total of 20 mice were vaccinated with ZRS05, with two different escalating dose regimens: 30µg (10µg of antigen, Group 1) and 60µg (20µg of antigen, Group 2). As controls, ten mice where vaccinated with antigen free formulation (Group 3) or left untreated (Group 4) (Table 1). Spleens were harvested seven days post-boost, and IFN-γ ELISpot was performed to measure the presence of SARS-CoV-2- and MERS-reactive CD4 and CD8 T cells. Despite induction of antibody responses, CD4 and CD8 T cell responses to both SARS-CoV-2 and MERS-specific antigens were not detected for any dose regimen (Supplementary Figure 1). As expected, the positive control stimulation with Concanavalin A (ConA) showed a robust increase in the number of IFN-γ SFUs, confirming the functionality of the ELISpot assay (Supplementary Figure 1).

## DISCUSSION

By developing a vaccine that also targets zoonotic elements, we can expand the immunogenicity cross-reaction to not only address emerging SARS-CoV-2 variants, but also address successful jumps from zoonotic virus reservoirs to humans and future epidemics and pandemics. In our vaccine development, we combined sequences from SARS associated with bats. Our zoonotic coronaviruses multivalent antigen strategy is based on the principle of a xenogeneic vaccine, which leverages components from one species to provoke an immune response in another species. In this context, the vaccine uses a protein associated with a pathogen that infects a different species but is structurally similar to the pathogen affecting humans. The key notion behind this strategy is the concept that numerous pathogens infecting varied species often share similar components or mechanisms. Utilizing these shared parts from a different species in the vaccine formulation could circumvent potential issues of immune tolerance, ensuring a more robust immune response.

Given that scalability, affordability, and acceptability are major constrains for vaccine programme implementation in low-middle income countries, the technology used for new vaccine candidates should consider these factors. Our vaccine strategy used a eukaryotic expression system to produce protein-based recombinant antigens vaccine strategy for its safety and proven track record in the vaccine field^26–31^. Given that protein antigens are usually poorly immunogenic on their own, we explored a range of adjuvants. We initially investigated the well-established alum adjuvant, in combination with either MPLA or CpG, to ensure a broad immune response to the vaccine. Alum is the most widely used adjuvant in vaccines for its safety, efficacy, and long history. MPLA and CpG are more recent additions to the vaccine adjuvants portfolio, and have high efficacies, but are difficult to scale and are costly. During a pandemic, such as the one with COVID-19, cost and supply issues were encountered with these three adjuvants. We therefore turned our attention to the Squalene-in-Water Emulsion (SWE) adjuvant, developed by a Seppic (Air Liquide Group) and the Vaccine Formulation Institute (VFI) partnership. Sepivac SWE™ has been made available in open access to the vaccine community by VFI^32–34^.

We found a pan-variant antibody response against SARS-CoV-2 and related coronaviruses when the Spike S1 subunit of the original Wuhan SARS-CoV-2 was formulated together with the S1 subunits of SARS-CoV-1 (the bat zoonotic element) and SARS-CoV-2 Alpha or Beta or Gamma variants and adjuvanted with alum-MPLA or alum-CpG. We elicited a robust increase in antibody levels against all SARS antigens tested (Wuhan, Bat, Alpha, Beta, Gamma, Delta, Kappa, and Omicron). Alum-MPLA and alum-CpG adjuvant combinations were equally effective, with no difference in antibody levels.

Finally, in our first squalene-based formulation (ZRS04), we observed an effective pan-variant antibody response against SARS-CoV and SARS-CoV-2 variants but no cross-reactivity to MERS. This is corrected in our final design, ZRS05, with the addition of the relevant MERS antigen while still conserving the SARS-CoV-2 cross-reactivity to relevant variants. Unfortunately, we did not detect SARS-CoV-2 nor MERS T-cell responses in mice immunized with the combined ZRS05 squalene-based formulation. This may be due to several factors, including the adjuvant used, the timing of sample collection, and the magnitude of the T-cell response. Further investigations with different adjuvants and larger sample sizes are warranted to confirm these findings and provide a comprehensive understanding of the T-cell response to ZRS05.

## Author Contributions

JAGP, DFR, AAP, SM, CMS, VL, BW conceived the study. AC, MDY, SE, MOD carried out experiments. SM, CMS, MM, MDY, MOD carried out T cell experiments. PAD, JW carried out antigen studies. GW, SM formulation and adjuvant work. JAGP, DFR, AAP, SM, CMS, AC, MDY wrote the manuscript with input from all authors.

## Acknowledgments

We are thankful to Prof. Mala Maini at the UCL Institute of Immunity and Transplantation and Ms. Marina Santilli UCL Business for all help and support provided.

